# Expression variation and covariation impair analog and enable binary signaling control

**DOI:** 10.1101/244236

**Authors:** Kyle M. Kovary, Brooks Taylor, Michael L. Zhao, Mary N. Teruel

**Affiliations:** Dept. of Chemical and Systems Biology; Stanford University; Stanford, CA, 94025

**Author notes:** **Corresponding Author:** Mary Teruel.

**Keywords:** single-cell proteomics, cell signaling, single-cell biology, cellular heterogeneity, MAPK/ERK signaling

## Abstract

Due to noise in the synthesis and degradation of proteins, the concentrations of individual vertebrate signaling proteins were estimated to vary with a coefficient of variation (CV) of approximately 25% between cells. This high variation enables population-level regulation of cell functions but abolishes accurate single-cell signal transmission. Here we measure cell-to-cell variability of relative protein abundance using quantitative proteomics of individual Xenopus laevis eggs and cultured human cells and show that variation is typically much lower, in the range of 5–15%, compatible with accurate single-cell transmission. Furthermore, we show that MEK and ERK expression covary which improves controllability of the fraction of cells that activate bimodal ERK signaling, arguing that covariation has a role in facilitating population-level control of binary cell-fate decisions. Together, our experimental and model data argues for a control principle whereby low covariation limits signaling noise for accurate control analog single-cell signaling. In contrast, increased covariation widens the stimulus-range over which external inputs can regulate binary cell activation, thereby enabling accurate control of the fraction of activated cells at the population level.

## Introduction

Vertebrate signaling has been shown to control both binary and analog outputs. Here we use the term binary if the output is bimodal and the term analog if the output signal changes in parallel with the input signal without bifurcations during the transmission. Examples of binary signaling decisions include the commitment to start the cell cycle (Cappell et al., 2016), cell differentiation (Ahrends et al., 2014; Chang et al., 2008; Jukam and Desplan, 2010), apoptosis (Spencer et al., 2009), action potentials (Hodgkin and Huxley, 1952) and the explosive secretory response of mast cells when encountering an antigen (Hide et al., 1993). Effective analog signaling in individual cells has been observed, for example, in the visual transduction system where the number of absorbed photons proportionally increases electric outputs in cone cells (Arshavsky et al., 2002), in single-cell IP3 and Ca2+ regulation by GPCR’s (Nash et al., 2001), as well as for CD-8 (Tkach et al., 2014) and IL-2 signaling (Feinerman et al., 2008) in T-cells. Analog signaling is also needed to accurately regulate the timing or duration of intermediate cell processes such as in the cell cycle where the time between the start of S-phase to mitosis has only small variation between individual cells (Spencer et al., 2013). Such precise regulation of durations requires low noise in the signaling steps before mitosis (Kar et al., 2009). Together, these examples suggest that accurate analog signaling is important for graded control of cell outputs in single cells as well as for accurate internal timing.

A main motivation for our study were the high levels of protein expression variation that have been reported in vertebrate cells with coefficient of variations (CVs) of approximately 25% (Gaudet et al., 2012; Sigal et al., 2006; Spencer et al., 2009). Such high levels of expression variation are beneficial for binary signaling which is often regulated at the population-rather than single cell-level. In population-based signaling, a goal of organisms is to use different levels of input to regulate the fraction of cells in a population that make a binary decision such as whether to proliferate, differentiate or secrete. For input stimuli to control which fraction of cells are activated, high noise in signaling is needed between cells in the population such that individual cells have different sensitivities to input stimuli (Ahrends et al., 2014; Eldar and Elowitz, 2010; Kalmar et al., 2009; Raj and van Oudenaarden, 2008; Süel et al., 2007). However, the same high noise needed to control population-level signaling does not have any benefit for analog signaling and just serves to degrade signal transmission. These different demands on noise for analog and binary signaling suggest that there is a trade-off for noise between population-level and single-cell signaling (Suderman et al., 2017). Nevertheless, the reported high levels of expression variation and signaling noise in mammalian cells (Cheong et al., 2011; Gaudet et al., 2012; Selimkhanov et al., 2014; Sigal et al., 2006) raise the question of how noise in a signaling system can be low enough for analog signaling accuracy. It also remained unclear how the different potential internal noise sources could generate optimal conditions for analog single-cell versus binary population-level signaling.

Here we measure cell-to-cell variation in the relative abundance of pathway components to understand the limits of analog and binary signaling accuracy. We also investigated the role of covariation of pathway components as we considered that covariation may exacerbate the analog signaling problem and/or enable the control of population-level binary signaling. We considered that previous estimates of cell-to-cell variation in protein expression might be too high due to experimental challenges in accurately measuring small differences in protein abundance between cells and accounting for “hidden variables” such as differences in cell size and cell cycle state (Symmons and Raj, 2016). To determine lower limits of protein variation, we developed single-cell quantitative proteomics methods in single *Xenopus laevis* eggs and employed quantitative normalization of cultured human cells to accurately measure variations in protein abundance normalized by protein mass. We found that cell-to-cell variation in relative protein abundance is much lower than expected, with CVs of between 5% and 15%, suggesting that expression variation is less limiting than currently believed and is compatible with accurate analog signal-transmission. Furthermore, our simulations show that these experimentally-observed low levels of expression variation pose a challenge for cells to accurately control population-level decisions. A solution to this problem was revealed by experiments which showed significant covariation between the single-cell expression of two sequential signaling components, MEK and ERK, and a correlation between MEK and ERK expression levels and whether or not ERK is activated. Our modeling showed that such increased covariation - which increases the overall noise in the signaling pathway – allows populations of cells to control the fraction of cells that activate ERK over a wider range of input stimuli, providing a potential role for covariation of signaling components as a general strategy for populations of cells to more accurately control binary cell fate decisions. Finally, our findings of covariation and low variation of pathway components led us to develop a metric to describe how systems can use opposing strategies to accurately control single-cell analog and population-level binary signal transmission by using different numbers of regulatory components, levels of expression variation, and degrees of covariation.

## Results and Discussion

### Computational simulations using reported levels of expression variation show a dramatic loss of analog single-cell transmission accuracy

Our study was motivated by the reported high levels of expression variation and the detrimental impact that this source of noise may have on analog single-cell signaling, especially since signaling pathways typically have multiple components which necessarily results in an even higher cumulative signaling noise. To define the general control problem of how expression variation increases overall signaling noise and limits signaling output accuracy, we carried out simulations by applying a relative fold-change in input signal (R) to a signaling pathway and stochastically varying the expression of pathway components for each simulation. To determine how accurately a multi-step signaling pathway can transmit a relative input stimulus (R) to an analog output (A*), we modeled the signaling pathway as shown in Figure 1A. Specifically, we used a 5-step model where a relative change in input R acts through four intermediate steps, possibly reflecting a kinase cascade with counteracting phosphatases, to generate corresponding changes in the output A*. The regulation of these steps can be at the level of activity or localization of pathway components. We considered 5 steps with 10 variable regulators to be a typical signaling pathway since it has been shown that step numbers in signaling pathways can range from very few in visual signal transduction (Stryer, 1991) to over 10 steps in the growth-factor control of ERK kinase and cell cycle entry (Johnson and Lapadat, 2002). In our simulations, each of the parameters represents a regulatory protein that activates or inactivates one of the pathway steps. We assumed that each of these components has “expression variation”, meaning that each of their protein concentrations varies between individual cells with a standard deviation divided by the mean (CV). We simulated this expression variation by multiplying a log-normal stochastic noise term of either 5%, 10%, or 25% (standard deviation over the mean) to each parameter in the model (Ahrends et al., 2014). As is apparent in the top plots in Figure 1B for a CV of 5%, the signaling responses of cells to 3-fold (red) and 9-fold (blue) increases in the input stimulus, R, can be readily distinguished from the signaling responses of unstimulated cells (black traces). For a higher CV of 10%, the signaling responses to a 3-fold increase in R partially overlap with the unstimulated cell responses, and only the responses to a 9-fold increase in R can be unequivocally distinguished from unstimulated cell responses. For a CV of 25%, even responses to a 9-fold increase in input stimulus overlap with the responses of unstimulated cells, showing a dramatic loss in signaling accuracy.

**Figure 1.**
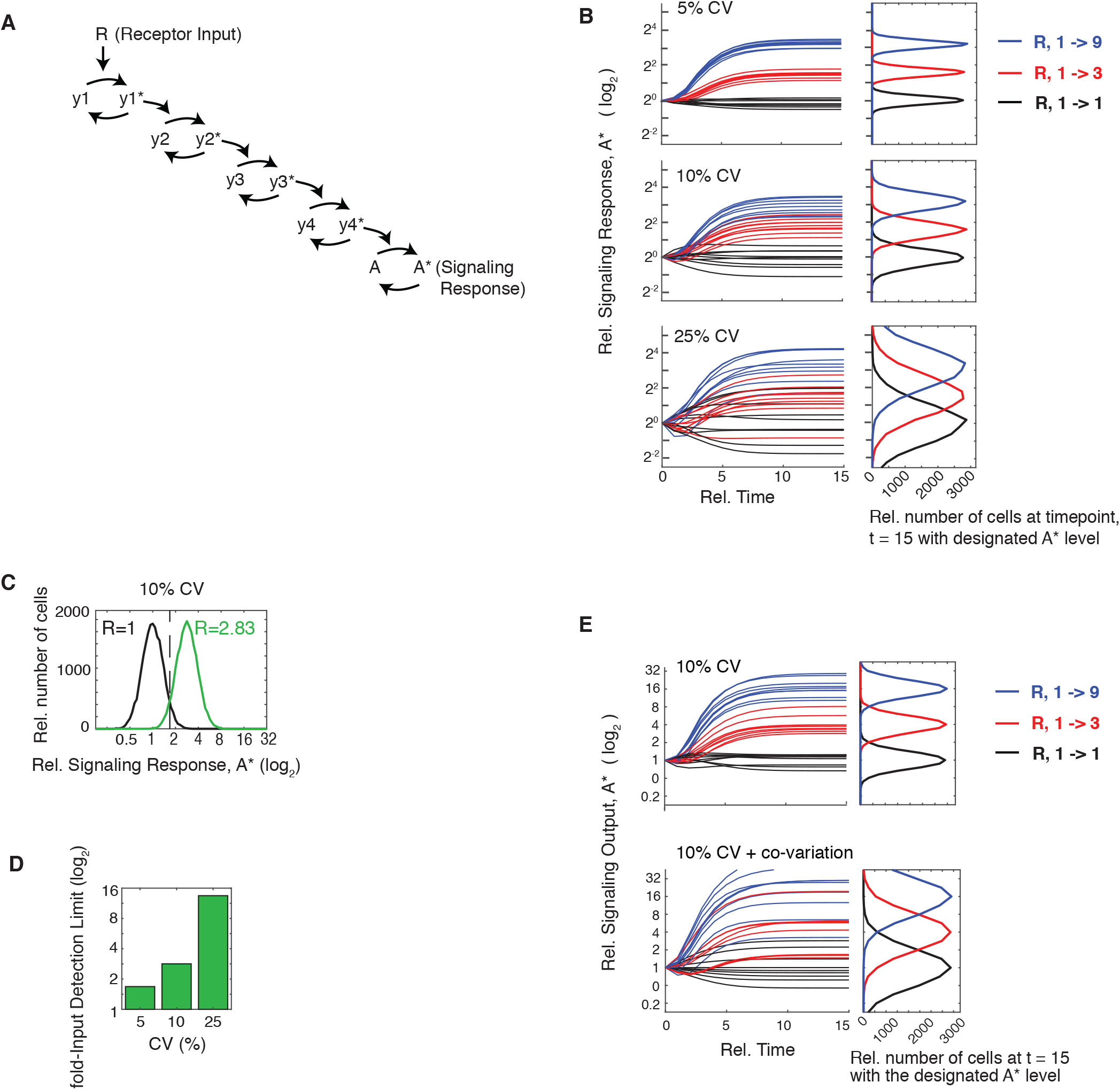
Computational simulations using reported levels of expression variation show a dramatic loss of analog single-cell transmission accuracy. (A) Schematic of a 5-step analog signaling pathway where the asterix(*) represents the activated form which is assumed in this model to be a small fraction of the total. (B) The timecourse plots show how relative 3- (red) and 9- (blue) fold input changes in R result in analog output responses with different degrees of noise. Random log-normal expression variation was added simultaneously to each pathway component. The accuracy of analog signal transmission is dramatically reduced as the coefficient of variations (CVs) increase from 5% (top),10% (middle), to 25% (bottom). (C) Top, Example of distributions of unstimulated (red) and stimulated (blue) cells at the fold-Input Detection Limit (fIDL). The fIDL represents the minimal stimulus needed to distinguish the output from stimulated from unstimulated cells with 95% accuracy, as marked by the vertical black dashed line. For the system in (A) with a 10% CV in each pathway component, the fIDL is 2.83. (D) Barplot comparing the fIDL values for the system in (A) with CVs of 5, 10 and 25%. (E) Simulation of the pathway model in (A) but now comparing the situation in which the pathway components are all uncorrelated with each other (top) with the situation in which the activating pathway components covary with each other and the de-activating pathway components covary with each other (bottom). The simulations in the right panel show that covariance of components of the same pathway would introduce a marked loss in signal transmission accuracy.

One way to overcome this dramatic loss in signaling accuracy due to expression variation of pathway components is to increase the input stimulus. We reasoned that we could use a fold-increase parameter to quantify the loss in signal accuracy. We thus defined a fold-Input Detection Limit (fIDL) as the minimal fold-stimulus needed to generate signaling responses that can, in 95% of cases, be distinguished from cell responses in unstimulated cells (see Methods for calculation). Figure 1C shows an example of how the fIDL is calculated by determining the fold-input stimulus that results in only a 5% overlap between the signaling output distributions (A*) of unstimulated and stimulated cells (blue and red histograms, respectively). In the case shown, an fIDL stimulus of 2.83 is needed to overcome the loss of signaling accuracy caused by having 10% expression variation in pathway components. We used a fold-Input Detection Limit instead of a commonly-used Mutual Information metric since Mutual Information between Input (R) and Output (A*) has a strong dependency on the dynamic range of the system output, while the fold-Input Detection Limit is largely independent of saturation (Figure EV1). As shown in the barplots in Figure 1D, increasing the CV of pathway components from 10% to 25% increases the fold-Input Detection Limit from 2.83 to 14, a stimulus requirement that is likely prohibitive for analog single-cell signal transmission. Our realization that fold-Input Detection Limits are very high for reported expression variation levels was a main motivation for our strategy below to accurately measure expression variation in order to understand if and how analog signaling in single cells is limited by this noise source.

We were also interested to determine if the expression of vertebrate proteins may covary since covariance has been shown to exist in a yeast regulatory pathway (Stewart-Ornstein et al., 2012). We considered that if proteins within a signaling pathway would covary, the overall noise in the output response would increase. To illustrate what a maximal effect of covariance can be on a multi-step analog signaling pathway, we added covariation to the model shown in Figure 1A by making the positive regulators (e.g. kinases) covary together and also made the negative regulators (e.g. phosphatases) covary together. As shown in Figure 1E, covariance causes the error propagation to increase, and the overall variation of a signaling output is much higher compared to the case where variation of proteins in the same pathway are independent. Given this marked increase of the overall noise of the signaling response by covariation, one would expect that covariation between components of the same signaling pathway should generally be avoided to allow for accurate analog signaling.

### Development of a method to accurately measure the relative abundance of tens of proteins in a single cell

To probe the lower limits of protein expression variation, we selected a system with a need for analog single-cell signaling that was also suitable for parallel proteomics analysis. We chose *Xenopus laevis* eggs for three reasons. First, previous studies showed that the timing of the cell cycle during early embryogenesis is very precise with an accuracy of ~5% (Tsai et al., 2014), suggesting that the Xenopus system must have accurate analog signaling to maintain such timing. Second, eggs do not grow in size and have only minimal new synthesis and degradation of mRNA, two features which we thought would reduce experimental and technical variation that was unrelated to cell-internal control of protein concentration. Third, *Xenopus laevis* eggs are well suited for single-cell proteomics analysis due to their large size (Ferrell, 1999), allowing us to very sensitively measure and compare relative abundances of many proteins simultaneously in the same cell.

To accurately compare the abundance of tens of endogenous proteins in parallel in single cells, we used a low-noise quantitative mass spectrometry method (SRM-MS) (Abell et al., 2011; Ahrends et al., 2014; Picotti and Aebersold, 2012). Cytoplasmic proteins were extracted from eggs and subjected to trypsin digestion and phosphatase treatment before undergoing targeted quantification on a triple quadrupole mass spectrometer. Heavy isotope-labeled reference peptides were spiked in proportionately to a measured total protein concentration, and the ratio of the light (endogenous) peptide to the heavy (synthetic) peptide was used as a readout of relative protein abundance. Small calibration errors were further corrected for during the analysis using the median of 22 normalized peptide intensities as a correction factor similar to previous studies (Abell et al., 2011; Feng and Picotti, 2016; Ludwig et al., 2012). We measured relative protein abundance (abundance over total protein mass) as a measure of protein concentration since reaction rates and signaling processes depend on the concentration rather than abundance of proteins (Padovan-Merhar et al., 2015).

We first validated our method using bulk cell analysis at different timepoints during the first cell cycle which can be initiated by addition of calcium ionophore and takes approximately 90 minutes to complete (Rankin and Kirschner, 1997). We measured the abundances of a set of 26 proteins that we selected to include known regulators of signaling and cell cycle progression, as well as several control proteins (Figure 2A; Table EV1). Time course analysis over the first cell cycle further showed that we could observe the expected cycling behavior of Cyclin A and Cyclin B (Figure 2B). We next showed that we could measure timecourses of relative protein abundances in single cells by carrying out measurements at 5 time points with 5 eggs each (Figure 2C). Except for a few known cell-cycle regulated genes, Cyclin A, Cyclin B, Cdc6 and Emi1, all of the measured proteins changed their abundance on average less than a few percent during the first egg cell cycle (Peshkin et al., 2015). The constant average level of many of these signaling and cell cycle proteins can in part be explained by only minimal mRNA synthesis during early *Xenopus laevis* cell cycles (Krauchunas and Wolfner, 2013).

**Figure 2.**
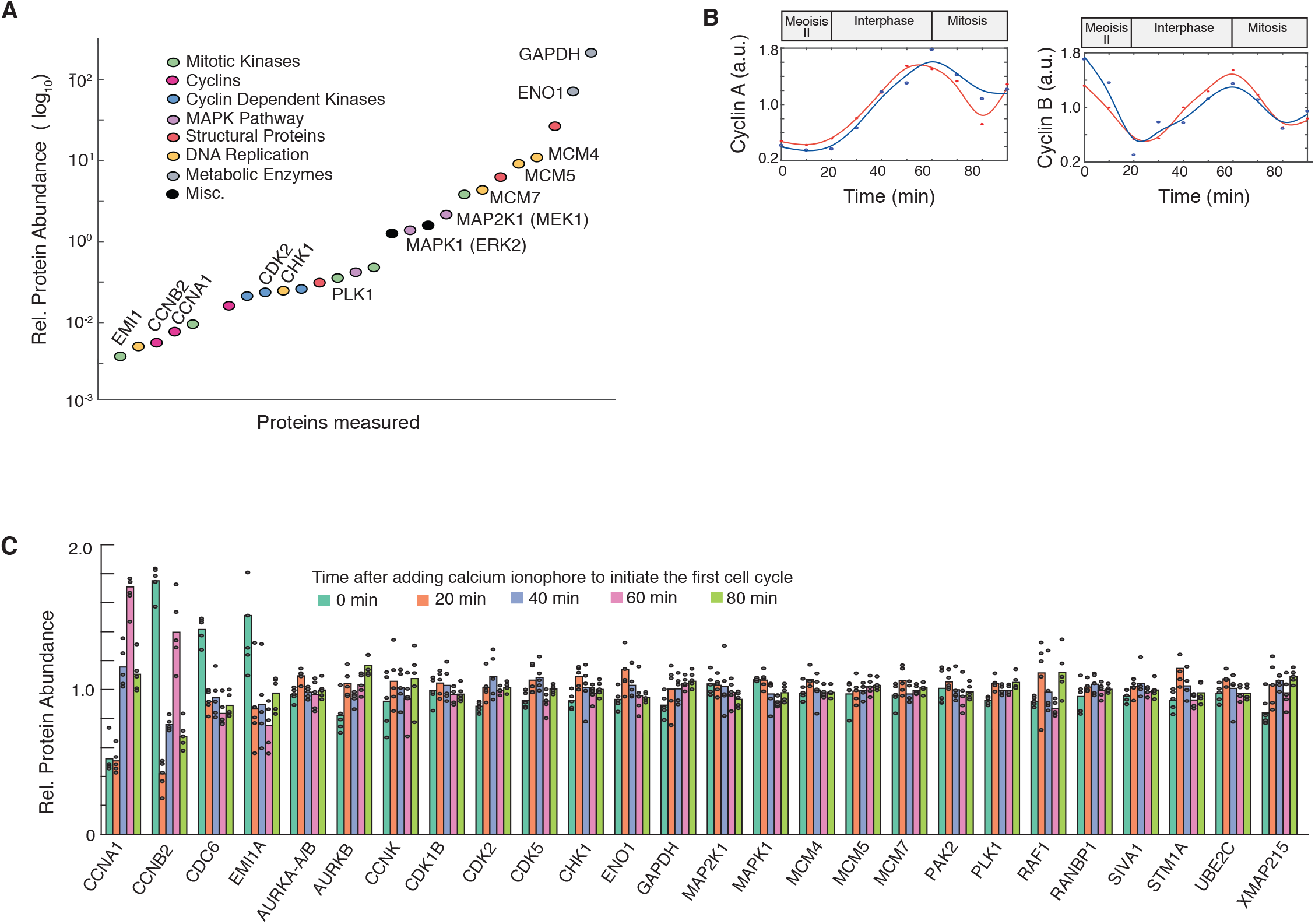
Development of a method to quantitatively measure relative abundances of tens of endogenous proteins in parallel in single Xenopus eggs. (A) Comparison of protein abundance of a set of cell cycle, signaling and control proteins in Xenopus eggs. Abundance measurements are based on SRM-MS mass spectrometry measurements of the combined cell extracts from 10 eggs per timepoint. Quantitation of relative protein abundance was carried out by adding heavy isotope-labelled reference peptides to the egg extracts. (B) Time course analysis of changes in Cyclin A and Cyclin B levels during the first Xenopus cell cycle. (C) 5 individual eggs were collected at 5 timepoints: 0, 20, 40, 60 and 80 minutes after the addition of calcium ionophore. To minimize variability due to sample handling and instrument sources, the 25 individual eggs were prepared for mass spectrometry analysis at the same time and were then analyzed in sequential runs on the same mass spectrometer. Barplot shows relative abundance changes of the 26 proteins shown in (A) tracked through the first egg cell cycle.

### Low variation in the relative abundance of proteins explains how cells can accurately control analog single-cell functions

We next focused on analyzing the extent to which protein abundance varies between single cells. We first analyzed the set of 25 individual eggs from Figure 2C and determined the variation of each protein in each of the batches of 5 eggs collected at each of the 5 time points (Figure 3A, left). Markedly, all CVs were much lower than expected with the median CV across all proteins and timepoints being only 7% (Figure 3A, histogram in right panel). To independently verify these low variation measurements, we collected and analyzed a larger set of 120 individual eggs: 60 eggs collected at 60 and at 80-minutes after activation. To test for reproducibility of the measured variation, we divided the 60 eggs at each timepoint into batches and carried out a variation analysis (Figure 3B). Bootstrapping analysis showed similar low variation (Figure EV2). As further validation, the variations measured in the two independent experiments were similar to each other (Figure 3C). We also noted that most of the proteins that have high cell-to-cell variation (marked as red circles in Figure 3C) also change their abundance during the cell cycle (Figure 2C) suggesting that high CVs reflect proteins whose abundances are actively regulated. Thus, our finding of low CVs answers the question raised in Figures 1A-C how cells can accurately control analog single-cell signaling outputs. Since expression variation can be as low as 5–10%, this main source of signaling noise is compatible with accurate single-cell signaling and timing control. Such low variation may also permit accurate timing in the *Xenopus laevis* embryonic cell cycle, which has been measured to be on the order of +/- 5% between eggs (Tsai et al., 2014),

**Figure 3.**
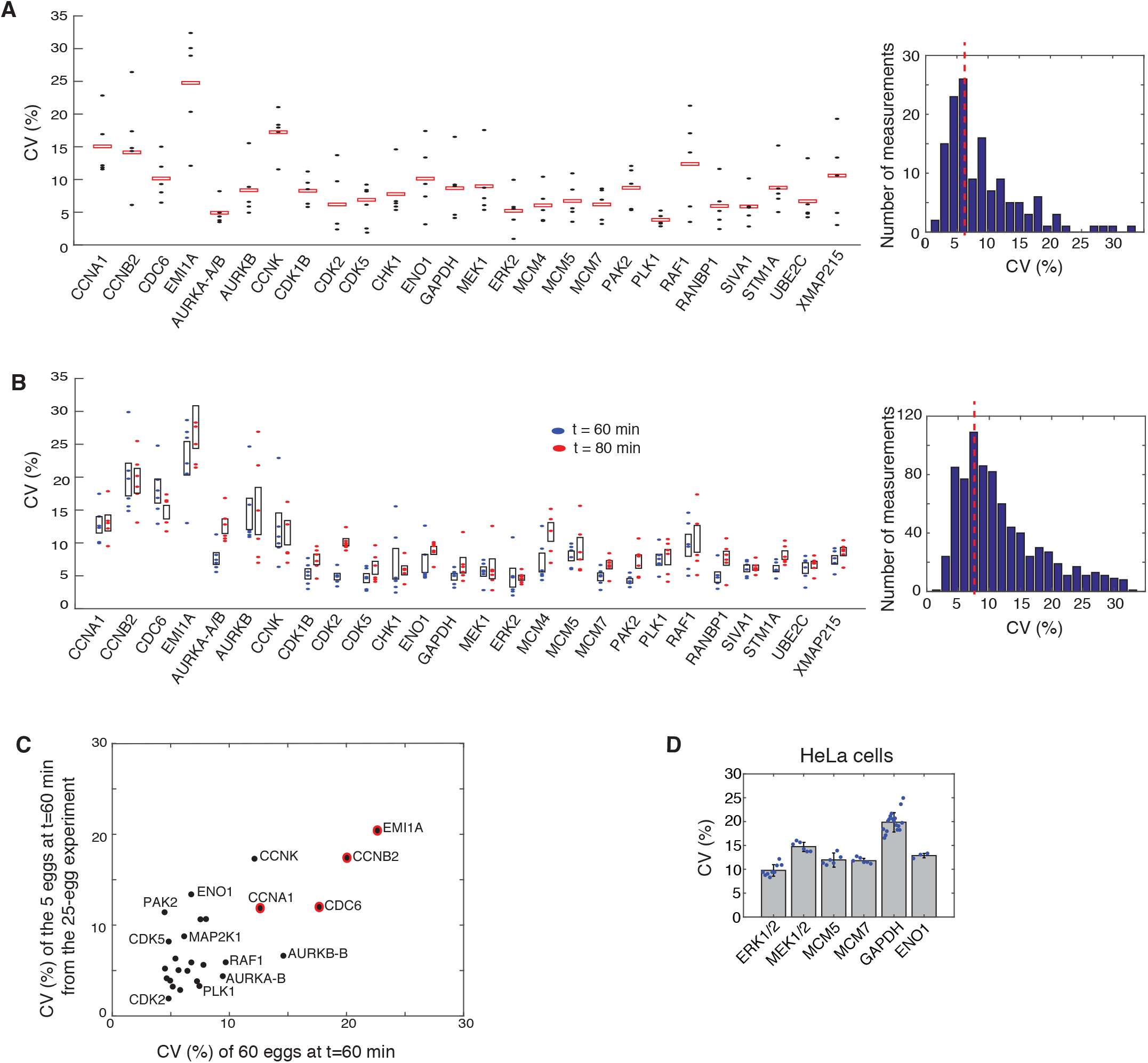
Single cell variation in relative protein abundance is typically 5–10% in Xenopus eggs. **(A)** Variation analysis of the relative abundance data from Figure 2C. Each blue point represents the coefficient of variation (CV) of the relative abundance of a protein between 5 individual eggs in a batch collected at the same time point. Red boxes mark the mean CV of 5 batches collected either at 0, 20, 40, 60 minutes after cell-cycle activation. (Right) A histogram of the measured CVs for all 26 proteins at 5 timepoints shows that CVs typically range from 5–10% with a mean CV of 7%. **(B)** Variation analysis of a second independent set of 120 eggs. 60 individual eggs were collected at 60 (blue) and at 80 (red) minutes after the addition of calcium ionophore. The 60 eggs at each time point were divided into 6 batches of 10 eggs analyzed sequentially on the mass spectrometer to minimize technical variation. The CV of the relative abundance of each protein between 10 individual eggs in a batch was calculated and plotted as filled blue and red ovals. The black boxes mark the 25th to 75th percentile of the 6 batch measurements for each protein at either the 60 or 80 minute timepoint. (Right) A histogram of the 12 CV measurements (6 CVs at 2 timepoints) for the 26 measured proteins shows the mean CV is 9%. **(C)** Control scatter plot of the CVs of the 26 measured proteins shows that measured protein expression variation is similar between 2 independent experiments, the 25-egg experiment shown in (A) and the 60-egg, 60-min experiment shown in (B). Red circles indicate proteins that have both high CV and change their abundance during the cell cycle. **(D)** CVs for a set of human homologs in HeLa cells. Immunocytochemistry was performed on cells plated in 96-well wells (representative images are shown in EV Figure 3). Each blue dot represents the CV calculated from the ~5000 cells in the respective well. Each barplot shows the mean CV of 3–12 wells. Error bars show standard deviation of the wells for that condition. Data shown is representative of 3 independent experiments.

We would like to note that the biological variation might be for some proteins even lower than we were able to measure in these experiments. To test whether there is a lower limit for measuring variation, we carried out control experiments in which 30 individual eggs were lysed and mixed together to remove biological variability. This mixed lysate was then pipetted into 30 individual tubes, and the sample in each tube was prepared and analyzed separately by SRM mass spectrometry. The variation between these 30 individually prepared and analyzed aliquots of the same starting lysate were compared to obtain a measure of technical variation. As shown in Figure EV3, the technical variation is comparable to the lowest CV measurements we show in Figures 3A-C, suggesting that further technical improvements may reveal even lower biological variation.

Our analysis so far argues that expression variation can be much lower than previously assumed, which would enable accurate analog single-cell signaling as shown by how decreasing expression variation in Figure 1B allows for less overlap between unstimulated and stimulated cell responses. We next tested whether we would find the same low variation in protein expression in cultured human cells (HeLa cells) by carrying out immunocytochemistry experiments. To accurately measure relative protein abundances, we first gated for cells in the same G0/G1 cell cycle state by using Hoechst DNA stain measurements (2n-peak; Cappell et al., 2016). We further normalized the abundance of each protein to total protein mass in each cell. The latter was measured using an amine-reactive dye that stains all proteins in a cell (Kafri et al., 2013). Since total protein mass is proportional to cell volume (Grover et al., 2011), normalization by total protein mass can be used as a measure of protein concentration, analogous to the normalization we used in the single egg experiments. To minimize small illumination non-uniformities associated with imaging, we also confined our analysis to cells in the center area of images where the illumination and light collection is more uniform (see Methods). For comparison with the egg data, we measured corrected CVs for the relative abundances for ERK, MEK, MCM5, MCM7 as well as the control proteins GAPDH and ENO1 (Figure 3D). We validated that the antibody staining for ERK, MEK, MCM5 and MCM7 could be knocked down by the respective siRNAs (Figure EV5). The resulting CVs for relative protein abundance were in the 10–15% range, lower than typically reported mammalian protein CV values (Gaudet et al., 2012; Niepel et al., 2009; Sigal et al., 2006).

### Covariance between the relative abundances of pathway components facilitates the control of population-level binary signaling responses

We next determined if there was covariance between proteins. To measure covariance, we used the same 120-egg proteomic dataset shown in Figure 2B. As shown in Figure 3A, our correlation analysis uncovered several covarying regulatory proteins. For example, there was significant co-regulation between MCM5 and MCM7 (Figures 4A and 4B), which is expected since the loaded MCM Helicase is a multimeric complex whose members have been shown to be subject to non-exponential decay, likely due to free subunits being degraded before complexed subunits (Mcshane et al., 2016). Nevertheless, we were surprised to also find significant covariation between MEK (MAP2K1) and ERK (MAPK1) (Figures 4A and 4B) because such a covariance adds extra noise to the signaling pathway and would not be beneficial for accurate analog signal transmission. In order to test the statistical significance of the covariance, p-values were validated by multiple comparison testing using Bonferroni corrections (Table EV2), suggesting that both the MCM5/MCM7 and the MEK/ERK covariation are significant.

**Figure 4.**
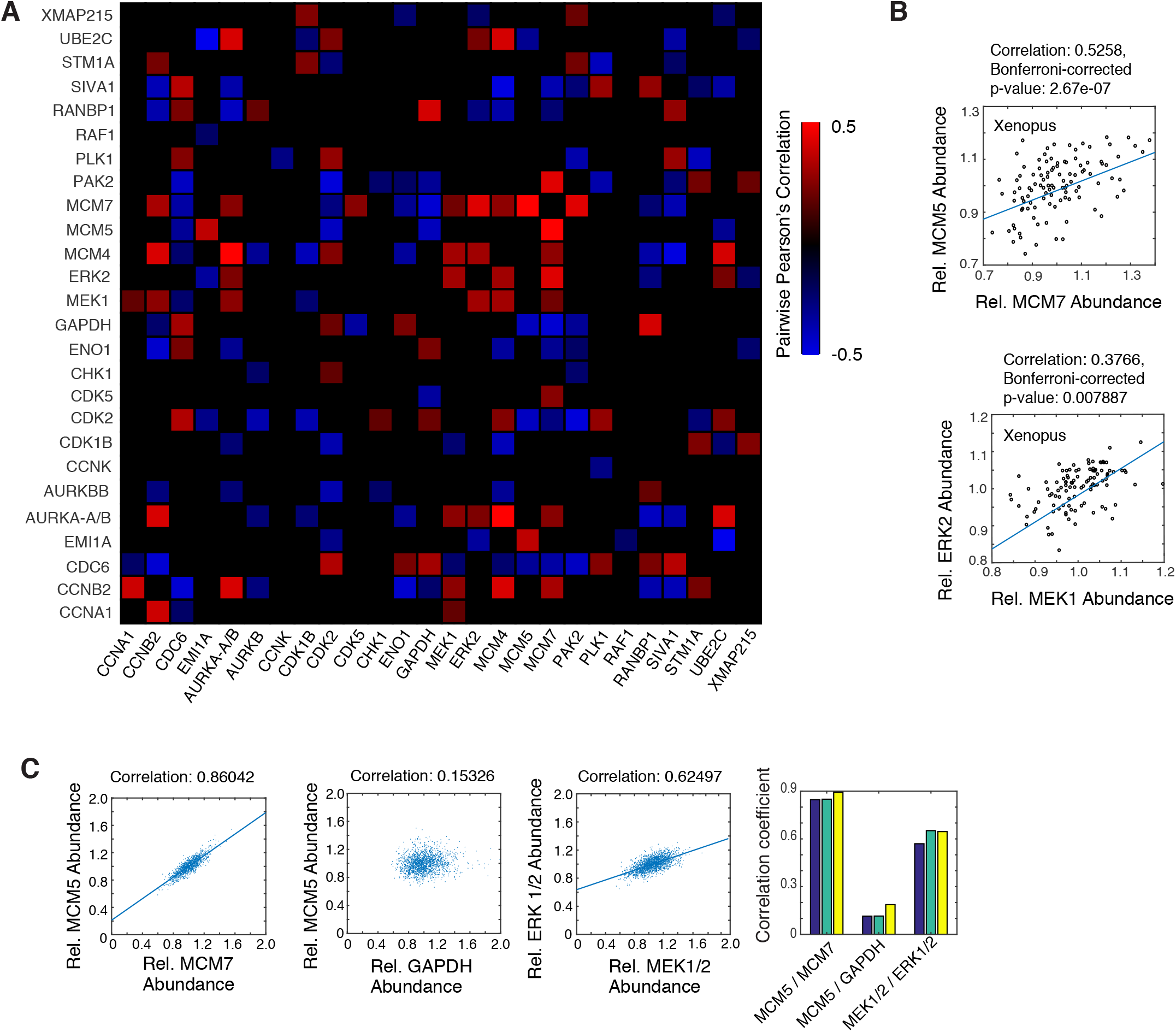
MEK and ERK expression covary in Xenopus eggs and cultured human cells. (A) Heatmap of Pearson correlation values between the respective proteins in Xenopus eggs. Twenty-six relative protein abundances were correlated pairwise in 120 single eggs. Only correlations with a p-value smaller than 0.05 are shown. P-values were validated by multiple comparison testing using Bonferoni corrections (Table EV2). (B) Two examples of pairwise correlations are shown between MCM5 and MCF7 and between MEK and ERK in Xenopus eggs. (C) Pairwise correlation analysis in HeLa cells, using MCM5 vs MCM7 as a positive control and MCM5 vs GAPDH as an uncorrelated control. Correlations between MEK and ERK concentrations are shown. Each scatter plot shows values from ~15000 cells. The bar graphs on the right shows correlation coefficients for 3 separate wells, containing ~5000 cells each, for the same 3 correlation pairs.

We next determined whether the covariances we observed in *Xenopus laevis* eggs are conserved in human cells. As shown in Figure 4C, we found a strong covariance between MCM5 and MCM7. siRNA-mediated depletion experiments confirmed that MCM5 and MCM7 likely co-stabilize each other as both levels are reduced upon knockdown of either MCM5 or MCF7 in HeLa cells (Figure EV5). While control experiments showed weak covariation between MCM5 and the control protein GAPDH, we once again found a significant covariation between MEK and ERK, similar to the covariance observed in *Xenopus laevis* eggs (Figure 4C). This co-regulation is likely due to shared upstream expression regulation, or indirect feedbacks, as siRNA-mediated depletion of MEK and ERK showed opposing effects on ERK and MEK expression, respectively (Figure EV5). The unexpected covariation found between MEK and ERK in both *Xenopus laevis* eggs and human cells made us consider whether it might be beneficial for a cell to have components of the same pathway covary, possibly in the context of binary cell activation that is often associated with MEK and ERK signaling pathways.

### Using modeling to understand the effect of variation and co-variation on controlling the fraction of cells in the population that respond to input stimulus (for binary single-cell signaling)

As mentioned in the introduction, previous studies showed that noise in signaling is beneficial for controlling the fraction of cells in a population that are activated over a range of input stimuli (Ahrends et al., 2014; Suderman et al., 2017). Since the MEK-ERK signaling pathway often controls binary proliferation and differentiation decisions, it was conceivable that the relevant output of the system could be at the population-level, reflecting the fraction of cells that would be activated or not. We carried out simulations to show how cell-to-cell variation and covariation in expression of pathway components would affect population-level regulation of binary cell fate decisions. As shown in the schematic in Figure 5A, we again used the model in Figure 1A but now assumed a last regulatory step whereby a cell with a y5* value above 10 would trigger a switch into an active state. However, if the output value y5* cell stays below a threshold of 10, the cell would remain inactive. This last step is denoted as B* versus B, reflecting the active and inactive binary output state, respectively. The results discussed here are largely independent of the value of this threshold (see Methods).

**Figure 5.**
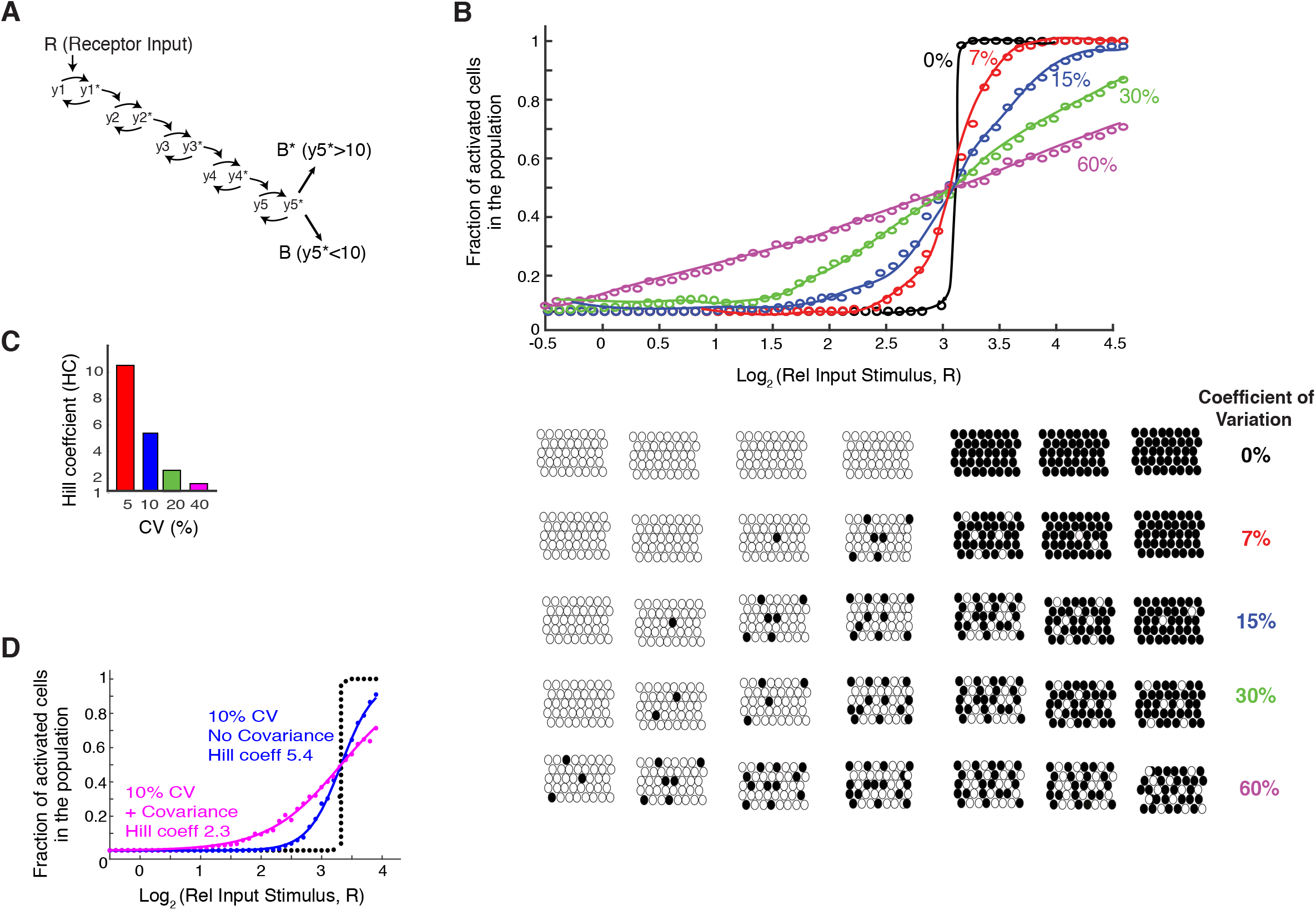
Using a general 5-step model to understand the effect of variation and co-variation on controlling the fraction of cells in the population that respond to input stimulus. (A) A binary output step was added to the model from Figure 1A. A threshold of 10 was used in each simulation to determine whether a cell was activated or not (y5*>10). (B) Plot of how increasing the CVs in expression of the pathway components in this binary model from 0% to 60% increases the range over which changes in the input stimuli can change the fraction of cells in the population which trigger the binary switch and become activated. (C) Hill coefficients (HC) were fit to the data in (B) to quantify the steepness in the curves. The steepness is an inverse measure of how wide the input range is that controls the output. (D) Same plot as in (B) but now comparing the output of the binary model if the pathway components vary randomly or covary with each other. The population response when uncorrelated CVs of 10% were applied to the pathway components is shown in blue. The magenta curve shows the population response when covariation was added to the model. To obtain a maximal effect, the CVs of 10% were applied to all positive and all negative regulators, respectively, such that the positive regulators covaried together and the negative regulators covaried together.

We used this binary model to determine the percentage of cells in a population that will switch into the active state for different fold-increases of input stimuli and levels of expression variation. As shown by the black circles in Figure 5B, if there is no expression variation of pathway components, all cells will abruptly switch from the inactive to active state for a very small increase of the input stimulus since all cells will either reach the output threshold of 10 or not for a given stimulus. As the CVs of relative abundances of pathway components increase, the percentage of cells in a population that switch from the inactive to active state can be controlled over a wider range of input stimuli, with there being a close to linear relationship in the five-step model between percent of cells activated and relative input stimulus amplitude at 40% CV of pathway components. This widening of the input stimulus control window can be quantified by fitting a Hill coefficient (HC) to the fractional activation data. The Hill coefficient measures how well the population-level output can be controlled by the input. The fitted Hill coefficients for systems with different amounts of protein expression variation are shown in the right bar plot of Figure 5C. A system with a smaller HC can be more accurately controlled over a wider-range of input levels which would be desirable in physiological settings where external hormone input stimuli may not be precise themselves. Another consideration to take into account is that physiological responses to hormone stimulation can typically be elicited over a 10-fold or greater range of relative hormone stimulus increases (R) (Atgié et al., 1997; Katakam et al., 2001; Kimura et al., 2007). Such a broad range of control requires a low Hill coefficient of approximately 1 (Figure 5B) which would require a system with high variation (approximately 40%) in the expression of pathway components or other sources of noise that vary the sensitivity of cells.

Given this need for low Hill coefficients to control population-level responses to physiological stimuli, we next determined whether covariation could be another source of noise that could lower the Hill coefficient and improve controllability. Such an increase in overall noise is needed as a system with 10% expression variation would not generate sufficient signaling noise for accurate population-control of binary signaling responses. Indeed, as shown in the simulations in Figure 5D, adding covariation to a regulatory system that has 10% variation of the pathway components improves the controllability of population-level binary responses by reducing the relationship between input stimulus and percentage of cells activated from a Hill coefficient of over 5 down to 2.3. Thus, our 5-step model demonstrates that a system with high covariation of signaling components enables population-level regulation of binary outputs over a broader range of signaling inputs. We next tested the validity of this conclusion computationally and experimentally by using an established model of the MAPK/ERK signaling system and by carrying out live-cell experiments to measure ERK activation in human cells.

### Using modeling and live-cell imaging to demonstrate the biological significance of MEK and ERK expression covariation in regulating bimodal ERK activity

We first explored the effects of covariation in the MEK-ERK pathway computationally using an established model of the MAPK pathway which uses 7 protein species: Ras, MEK, ERK, 4 phosphatases, and RasGTP as the input (Sturm et al, 2010). We varied the model parameters and tested the model over a range of RasGTP input doses. In the calculations shown, we either kept the concentration variations of ERK and MEK random or covaried them with each other (lognormal 15% CV), and also used random lognormal concentration variation for the other parameters/components in the model (lognormal 10% CV). With this added variation, the output of the model, phosphorylated ERK (pERK), which reflects ERK activity, becomes variable between cells and is bimodal for intermediate concentrations of EGF stimuli as shown by the timecourse traces in Figure 6A, as well as in the histograms in Figure 6B.

**Figure 6.**
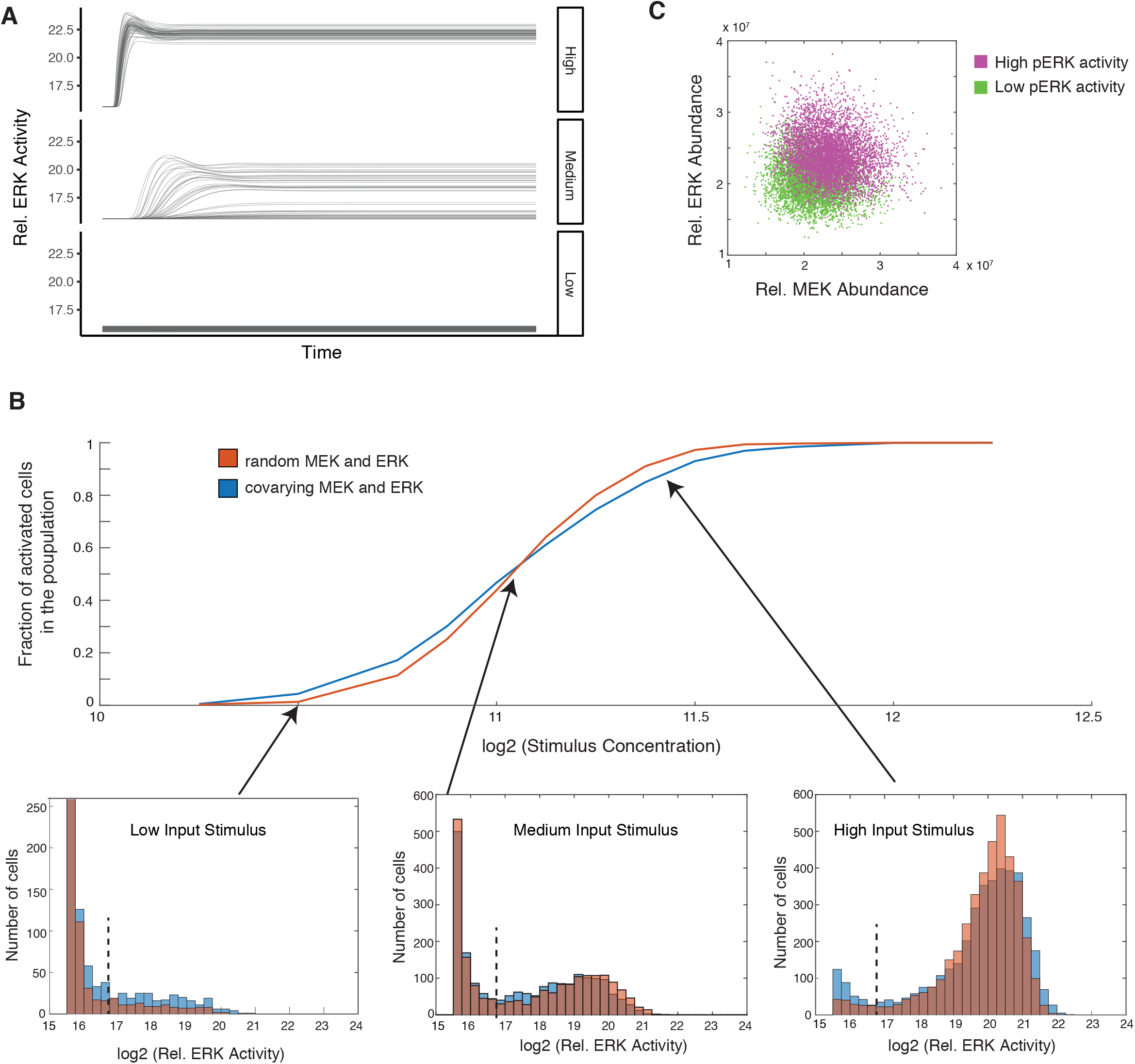
Simulations using an established MEK-ERK signaling model show that covariation between MEK and ERK expression widens the window over which input stimuli can control the fraction of cells that are activated in the population. (A) Timecourse output from an established MEK-ERK model (Sturm et al, 2012) in response to high, medium, and low concentrations of input (RasGTP) stimulus. Random lognormal noise with 15% CV was applied to MEK and ERK and with 10% CV to all other protein variables. (B) Comparison of the effect of covarying MEK and ERK concentrations. The orange curve and histograms show the results of simulations in which random lognormal noise with 15% CV was applied independently to the MEK and ERK concentrations. The blue curve and histograms shown the results of simulations in which MEK and ERK concentrations were made to covary by applying the same 15% CV lognormal noise term to both MEK and ERK in each cell/simulation. The bottom panels show examples of the distributions of outputs for three input stimuli. The dashed black line shows the threshold used to distinguish active from inactive cells. The shallower slope of the blue curve show that the fraction of activated cells can be regulated over a wider range of input stimuli if there is covariance between MEK and ERK. (C) Scatter plot of cells colored by whether they had high (magenta) or low (green) ERK activity at the end of the timecourse. Cells shown were stimulated with input doses between 2^10.5 to 2^12, a range which results in both active and inactive cells in the population as shown in (B).

The orange and blue curves in Figure 6B plot the fraction of cells in the cell population that go into a high pERK (activated) state in response to different doses of EGF stimulus in the model when MEK and ERK vary randomly (orange) or when MEK and ERK covary fully (100%) with each other (blue). When comparing the effect of random variation versus covariation of MEK and ERK concentrations, we find a small but significant broadening of the relationship between the stimulus intensity and the fraction of cells in the active state when there is covariation (Figure 6B, orange versus blue curves). This effect is proportionally reduced if the covariation is partial and would increase if more pathway components - in addition to MEK and ERK - would covary with each other.

The contribution of covariance can best be seen in the histograms shown at the bottom of Figure 6B. The histograms plot the number of cells with a particular ERK activity output in response to different doses of input stimuli. The black dotted line in each histogram marks the threshold above and below which cells are defined as having active or inactive ERK, respectively. The leftmost histogram shows that already for low input stimuli, the fraction of activated cells is more than doubled in the case of covariation compared to the case of random variation. The relatively higher fraction of activated cells in the case of covariation is caused by a widening of the range over which cells in a population are activated since for both the case of covariation or random variation, the model shows approximately the same number of cells - half of the population - are activated for the same intermediate input stimulus. The broadening of the range over which increasing input stimuli concentrations can control increasing fractions of activated cells can also be seen in the third histogram for high input stimuli: for both random variation and covariation, the high stimulus causes most cells to shift into the active state. However, twice as many cells still remain in the inactive state in the case of covariation compared to the random variation case. Together, this suggests that covariation broadens the window over which the fraction of activated cells in a cell population can be controlled by changes in input stimuli concentrations, which would be particularly important if organisms need to control the activation of small fractions of cells in a population such as to enable low rates of cell differentiation (Ahrends et al., 2014) or apoptosis (Spencer et al., 2009) in tissues. Finally, Figure 6C shows that cells in the population with high ERK activity have on average higher MEK and ERK levels compared to cells that have low ERK activity, arguing that the concentrations of MEK and ERK are limiting and thus matter in determining whether or not a cell will be activated.

In summary, these model calculations show that covariation of MEK and ERK expression can improve the population-level fractional control of ERK activation. For this model to apply to ERK activation in a particular cell type, a few conditions are critical: 1) The ERK signaling output should be bimodal or at least variable for intermediate stimuli in the same population. Variable outcomes for the same stimulus are needed if the goal of the signaling system is to control the fraction of cells in a population that is activated. 2) The expression of MEK and ERK should covary with each other in the particular cell system tested. Significant covariation is needed for it to contribute to broaden the input range over which cells can control the fraction of activated cells. 3) Finally, the structure of the ERK signaling model predicts that variation of expression levels of MEK and ERK proteins make a significant contribution to the variability of the ERK signaling output. To test whether this is indeed correct, the expression of MEK and ERK should on average be higher in cells with high ERK signaling compared to cells with low ERK signaling when analyzed in the same population of cells for the same intermediate input stimulus.

We tested whether these conditions are met in EGF-stimulated MCF10A cells. Specifically, we generated an MCF10A cell line expressing a FRET sensor of ERK activity to measure ERK activation in live cells (Albeck et al., 2013; Aoki et al., 2013). The FRET intensity of this sensor, EKAR-EV, was shown previously to faithfully report pERK levels in MCF10A cells (Yang et al., 2017). We used EGF to activate the pathway, and after 60 minutes, cells were fixed and stained with antibodies to measure the concentration of MEK and ERK, so that the pathway response could be related back to the relative level of the two proteins. An EGF titration was performed to test for bimodal ERK activation and to identify intermediate stimuli doses that induced heterogeneous responses (Figure 7A). We quantified the ERK activity in each timecourse by calculating integrated ERK activity as the area under the curve after EGF stimulation. As shown in Figure 7B, the integrated ERK activity values can be approximated as bimodal, allowing us to define cells as active or inactive using the indicated threshold (dotted vertical black line). From the histograms in Figure 7B, it is apparent that the fraction of activated cells in the cell population increases as the EGF concentration increases, and this relationship is more directly plotted in Figure 7C. Thus, in these human MCF10A cells, there is a wide range of input stimuli over which the fraction of the cell population that is activated can be controlled.

**Figure 7.**
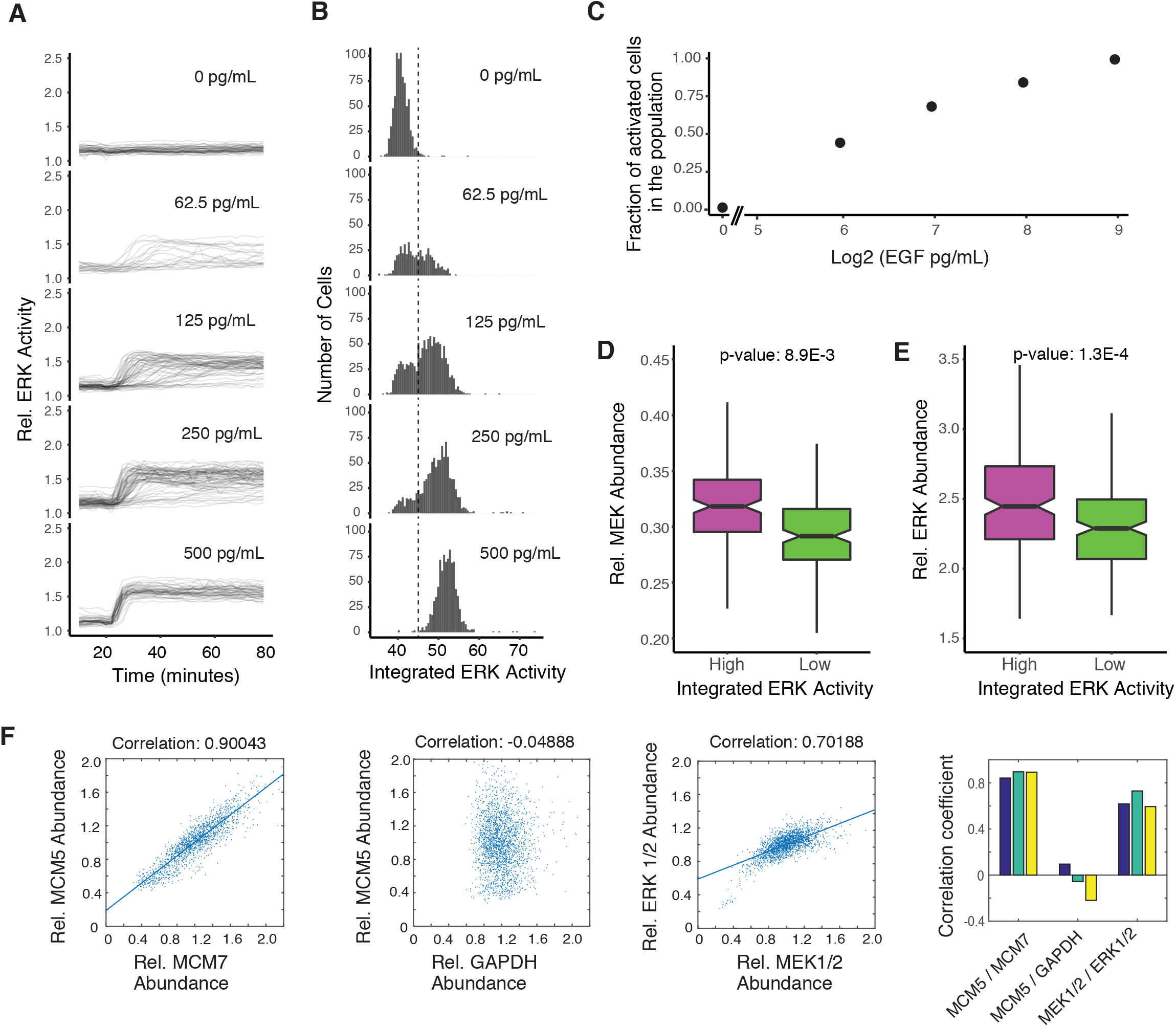
Single cell imaging experiments also show that covariation between MEK and ERK expression facilitates control of bimodal ERK activation. (A) MCF10A cells stably expressing the EKAR-EV FRET sensor were activated with varying levels of EGF after being serum starved for at least 48 hours. Cells were imaged every 2 minutes throughout the time course. The plots at EGF doses of 0, 62.5, 125, 250, and 500 pg/mL show timecourses from approximately 800, 520, 1200, 1000, and 900 individual cells, respectively. (B) Histograms showing the integrated ERK activity of individual cells from the experiment shown in (A). Integrated ERK activity was calculated for each timecourse as the area under the curve after the addition of EGF. The dashed line shows the threshold used to distinguish cells with active versus inactive ERK. (C) Plot showing fraction of activated cells (cells to the right of the threshold plotted in (B)) in response to different EGF concentrations. (D) Box and whisker plot of MEK concentration in cells with high (top 15%, magenta) or low (bottom 15%, green) integrated ERK activity. The high and low conditions represent 162 and 161 cells respectively, out of a total of 1073 cells. (E) Box and whisker plot of ERK concentration in cells with high (top 15%, magenta) or low (bottom 15%, green) integrated ERK activity. The high and low conditions represent 198 and 197 cells respectively, out of a total of 1316 cells. (D, E) The box plots show the distributions of protein concentrations in single cells with high versus low integrated ERK activity, where the bold line in the center of the notch represents the median, the ends of the notched box represent the first and third quartiles, the length of the upper whisker shows the largest point no more than 1.5 times the inter-quartile range (IQR or length of the box), the lower whisker represents the smallest point no more than 1.5 times the inter-quartile range, and the notches represent 1.58 * IQR / sqrt(n), which approximates the 95% confidence interval of the median. The non-overlapping notches between the high and low populations, as well as the low p-values, indicate that the differences between the two populations are significant. (F) Pairwise correlation analysis in MCF10A cells, using MCM5 vs MCM7 as a positive control and MCM5 vs GAPDH as an uncorrelated control. Correlations between MEK and ERK concentrations are shown. Each scatter plot shows values from ~15000 cells. The bar graphs on the right shows correlation coefficients for 3 separate wells, containing ~5000 cells each, for the same 3 correlation pairs.

The abundances of MEK and ERK in individual cells were measured at the end of the EGF timecourses by immunocytochemistry, and the values were normalized by an intracellular total protein stain following established protocols from (Kafri et al., 2013) in order to correct for cell volume and to obtain relative protein abundances. When we compared relative MEK and ERK abundances in cells with active or inactive ERK activity, we confirmed the prediction from Figure 6C that activated cells have on average higher MEK and ERK concentrations, and inactive cells have on average lower MEK and ERK concentrations (Figures 7D and 7E). Furthermore, we confirmed that MEK and ERK covary with each other in these cells (Figure 7F) with a correlation coefficient of 0.7. The covariation between MCM5 and MCM7 and lack of covariation between MCM5 and GAPDH are shown as controls Together, these modeling and experimental results support each other and show that covariation in the relative abundance of components in a signaling pathway can broaden the signaling variation of cells in a cell population and thereby facilitate the regulation of the fraction of activated cells at the population level.

### Constraints on accurate control of analog and binary signaling by expression variation and covariation

We used our experimentally-measured low CV values for relative protein abundances and our finding that covariation regulates binary signaling outputs, to explore the respective ranges of variation and covariation where single-cell and population-level signaling can be effectively controlled. As depicted in Figure 8A, we employed a modification of the model from Figure 1A to directly compare analog and binary signaling outcomes by assuming that the same pathway drives in one case an analog single-cell output (A*) and in the second case, binary cell activation if the output y5* reaches higher than a threshold of 10 (B*). We measure analog single-cell signaling accuracy by using fold-Input Detection Limits (fIDL), as defined in Figures 1C and 1D, and accurate controllability of population-level binary signaling by using Hill Coefficients (HC), as defined in Figure 5C. As discussed in Figure EV1, the fIDL parameter is a measure of analog signaling accuracy that is inversely related to mutual information but is less dependent on the dynamic range of the output, and the Hill coefficient is an inverse measure of the input range over which the population-level output can be controlled. The equations used to calculate the fold-Input Detection Limit and Hill Coefficient are shown on top of Figures 8B and 8C (see Methods for derivation).

**Figure 8.**
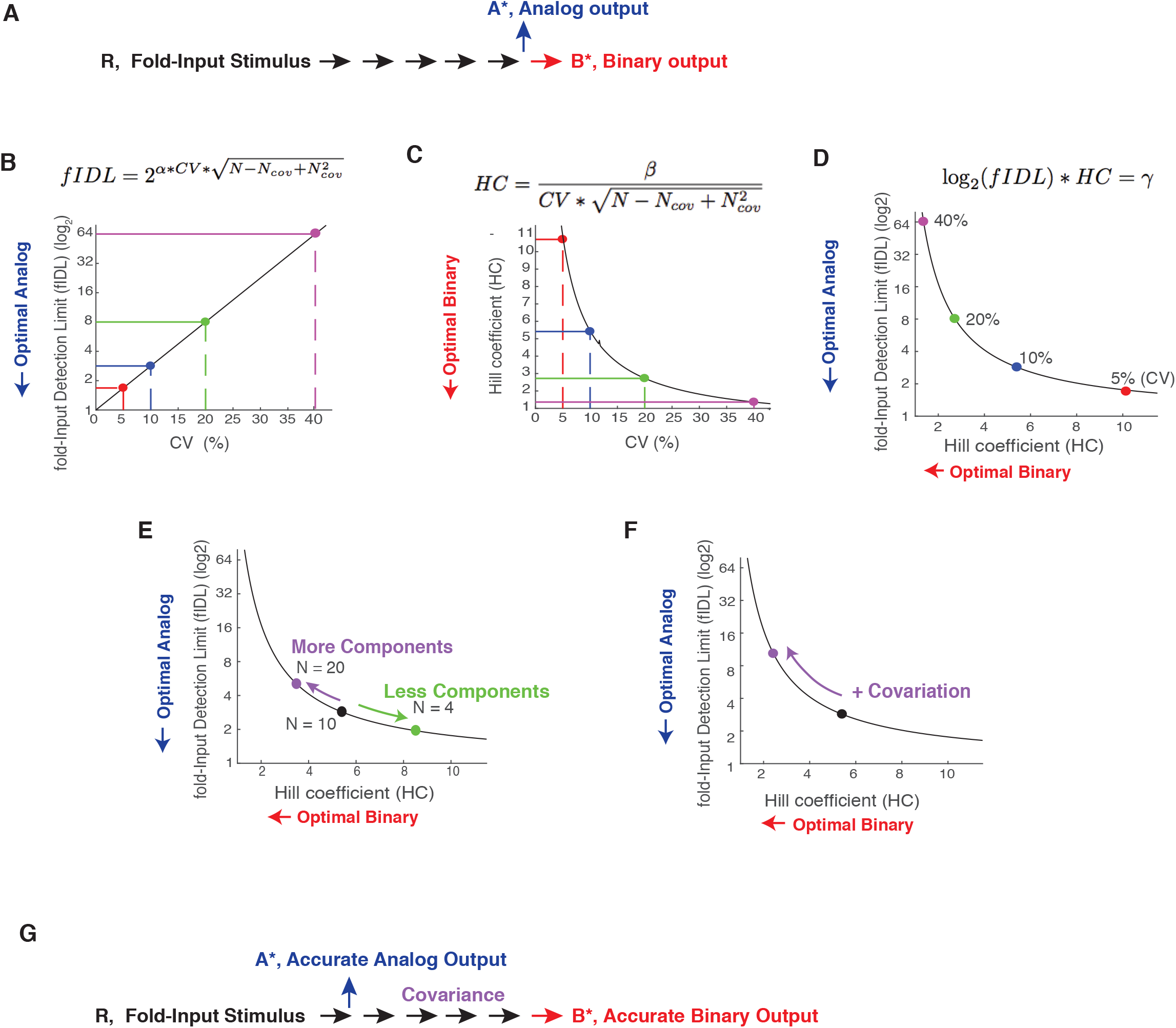
Competing demands on variation and covariation in the control of analog single-cell versus binary population-level signaling outputs. (A) Schematic of a signaling pathway that splits into an analog or binary output. (B-D) Quantification of the competing constraints on expression variation for accurate control of single-cell analog versus population-level binary signaling outputs. Plots showing the development of a metric that quantitatively relates expression variation, analog single-cell signaling accuracy, and binary signaling accuracy. (B) Relationship between expression variation and fIDL, highlighting the physiological range of 5, 10, and 20% expression variation. (C) Same for expression variation versus HC, also including 40% expression variation which is required for accurate control of population-level binary outputs. (D) Integration of both relationships into a single co-dependency curve relating optimal analog single-cell and binary population-level signaling. Terms in equations: CV: expression variation; N: Total number of pathway components; Ncov: Number of covariant pathway components; a = 3.29; b = 1.72; g = 6.29. (E) Increasing or decreasing the number of regulators in a pathway increases or decreases the overall noise in the pathway, respectively, and thus can be used as a way to more accurately control either binary population-level or analog single-cell functions, respectively. (F) Covariation between pathway components such as MEK and ERK is an effective means to increases overall noise in the pathway and thereby improve the controllability of binary population-level signaling responses while reducing single-cell analog signaling accuracy. A system with covariation can accurately control binary population level signaling without a need for 40% expression variation which is likely not common. (G) The analysis of (B)-(F) suggests that the same pathway components can only be shared between analog and binary signaling systems if the analog pathway branches off early after receptor stimulation. Covariance in a branch of a signaling pathway is an indication that the output is regulated by a binary output at the population-level.

As shown in Figures 8B and 8C, single-cell analog or population-level binary outputs can be optimally controlled if the fold-Input Detection Limit or Hill Coefficient, respectively, are small and close to 1. The conflicting constraint between the control of single-cell analog and population-level binary signaling by expression variation can be seen clearly by combining the two graphs in Figures 8B and 8C into a single competition curve (Figure 8D). Increasing the variation in the concentration of pathway components moves cells along this curve from optimal conditions for analog single-cell signaling (CV of 5%, right bottom) towards optimal conditions for control of binary population-level signaling (CV of 40%, left top) with the curve staying far away from the origin at the left bottom where analog and binary signaling would both be accurate. Thus, the same signaling system with a CV of 5% that has optimal analog single-cell accuracy loses its ability to accurately control binary population-level outputs. Similarly, a system with a CV of 40% that is optimal for controlling binary population-level outputs loses its ability to accurately control analog single-cell signaling. Thus, cells cannot have a shared pathway that controls accurate analog single-cell signaling outputs and also accurately controls binary population-level signaling outputs.

As shown in the Figures 8C and 8D, as well as in Figure 5B, a CV of 40% or greater would be optimal for controlling population-level signaling outputs. However, our study and previous work by others suggests that such high CVs of protein concentrations are not common (Gaudet et al., 2012; Sigal et al., 2006), indicating that cells must use other mechanisms to generate the necessary high signaling noise to accurately control the fraction of activated cells for population-level binary outputs. We considered that changes in the number of pathway components as well as the covariance of pathway components are strategies to alter the overall signaling output noise. We used the fIDL versus HC co-dependency curve to first determine how changes in pathway component numbers control analog or binary signaling (Figure 8E). While our analysis so far assumed 10 regulatory elements, fewer or higher numbers of signaling steps are common in signaling systems. Notably, changing the number of signaling steps improves one signaling mode at the cost of the other. Fewer signaling steps move the system towards improved analog single-cell signal transmission and more signaling steps towards improved control of population-level binary outputs. To illustrate the effect of increasing or decreasing signaling steps with examples: since many signaling systems are complex with likely 20 or more regulators (Gaudet et al., 2012; Sturm et al., 2010), such complex systems must necessarily be mediating population-level signaling responses. In contrast, the visual signal transduction pathway in retinal cone cells, which transduces light intensity inputs proportionally into electrical outputs, has only a few main regulatory components (Arshavsky et al., 2002) which benefits the control of analog single-cell signaling responses.

Our modeling and experimental data in Figures 5–7 showed that a potent strategy to increase noise, without adding expression variation to individual components, is based on positive covariation between pathway components. Covariation can increase accurate binary signal transmission as we show in the case of the MEK-ERK signaling pathway. Indeed, Figure 8F shows that adding covariation moves cells away from a state where they can accurately perform analog single-cell signaling towards a state where they can accurately control the fraction of activated cells at the population-level. This suggests that covariation is a powerful strategy to improve the control of population-level binary cell functions without that the expression variation or number of pathway components themselves have to be increased. We also note that covariation can in some cases be used to improve rather than increase analog single-cell accuracy if directly opposing enzymes (e.g. a kinase and a phosphatase) covary with each other (Feinerman et al., 2008). Together, our analysis shows that cells have a versatile set of internal tools to control whether a signaling pathway can accurately control single-cell analog or population-level binary signaling by either changing the expression variation of individual components, the number of pathway components, or the covariation in expression between components. Furthermore, if pathways share components, these model calculations argue that analog signals have to minimize component numbers by branching out early in a pathway, while binary population-level signal responses would optimally go on for longer with pathway components covarying with each other (Figure 8G).

## Conclusions

Our study employed sensitive single-cell mass spectrometry of *Xenopus laevis* eggs and normalized single-cell immunofluorescence analysis in cultured mammalian cells to reveal a low variation in relative protein abundances in the 5–15% range, suggesting that expression variation is not prohibitively high for analog signal transmission in single cells as was often assumed in previous studies. Nevertheless, the relative abundance of components of signaling pathways still show significant variation, arguing that there is a lower biological limit for accurate analog single-cell signal transmission in cells. Our simulations in Figure 1A showed that low CV values of 5–10% can enable accurate analog signaling for input changes that are two-fold or greater. However, as shown in Figure 5, these low CV values are too low for accurate control of population-level binary signaling systems, arguing that other mechanisms besides just relative protein abundance variation must exist to increase signaling output noise. Our modeling showed that covariance of signaling components is an effective mechanism to increase output noise and thereby control binary cell responses at the population-level. We were able to experimentally confirm such a beneficial role of covariation in our study of MEK-ERK signaling in Figures 6 and 7. Nevertheless, our analysis in Figure 1E showed that covariation also has an opposing effect by harming the accuracy of analog single-cell signaling. Given these opposing roles of expression variation and covariation, our study answers a fundamental question of how cells can adapt these two core sources of signaling noise to generate signaling systems that can accurately control single-cell analog or population-level binary signaling responses. Finally, we used model simulations that show a control principle whereby noise contributions from number of pathway components and covariation can shift signaling systems with the same expression variation of pathway components either towards accurate analog single-cell or binary population-level signaling.

## MATERIALS AND METHODS

### Contact for Reagent and Resource Sharing

All reagents including cell lines and antibodies are listed in Table EV5. Further information and requests for reagents may be directed to, and will be fulfilled by the corresponding author, Dr. Mary N. Teruel.

### Xenopus Laevis Egg Collection and Activation

Xenopus egg extracts were prepared based on modifications of a previous protocol (Tsai et al., 2014). All of the animal protocols used in this manuscript were approved by the Stanford University Administrative Panel on Laboratory Animal Care. To induce egg laying, female *Xenopus laevis* were injected with human chorionic gonadotropin injection the night before each experiment. To collect the eggs, the frogs were subjected to pelvic massage, and the eggs were collected in 1X Marc’s Modified Ringer’s (MMR) buffer (1M NaCl, 20mM KCl, 10mM MgCl2, 20mM CaCl2, 50mM HEPES, pH 7.8). To remove the jelly coat from the eggs, they were placed in a solution of 2% cysteine in 1X MMR buffer for 4 minutes and gently agitated, after which they were washed 4 times with 1X MMR buffer. To activate the cell cycle, eggs were placed in a solution of 0.5ug/mL of calcium ionophore A23187 (Sigma) and 1X MMR buffer for 3 minutes, after which they were washed 4 times with 1X MMR buffer. Single eggs were collected at their respective time points and placed into 600uL tubes and snap frozen in liquid nitrogen before being stored at −80C.

### SRM Sample Prep

Single eggs were lysed mechanically by pipetting the egg in 100uL of lysis buffer (100mM NaCl, 25mM Tris pH 8.2, Complete EDTA-free protease inhibitor cocktail (Sigma)). The lysate was then placed in a 400uL natural polyethylene microcentrifuge tube (E&K Scientific 485050) and spun at 15,000g in a right angle centrifuge (Beckman Microfuge E) at 4 degC for 5 minutes. The lipid layer was removed by using a razor blade to cut the tube off just beneath it, and the cytoplasmic fraction was pipetted into a 1.5mL protein LoBind tube (Fisher Scientific 13–698–794), being careful to leave the yolk behind. To precipitate the proteins from the cytoplasmic fraction, 1mL of ice cold acetone was added to each sample and placed at −20 degC overnight.

To collect precipitated proteins, the samples were centrifuged at 18,000g for 20 minutes at 4 degC. Acetone was decanted and the protein pellets were resolubilized in 25uL of 8M urea. To fully solubilize the protein pellet, the samples were placed in a shaker for 1 hour at room temperature. The samples were then diluted to 2M urea with 50mM ammonium bicarbonate to a 100ul volume, after which protein concentration was measured in duplicate with a BCA assay by taking two 10ul aliquots of each sample. The proteins in the remaining 80ul of sample volume were reduced with 10mM TCEP and incubated for 30 minutes at 37degC, then alkylated with 15mM iodoacetamide and incubated in the dark at room temperature. Next the samples were diluted to 1M urea with 50mM ammonium bicarbonate, and heavy peptides (JPT SpikeTides) were added based on BCA assay results. Trypsin (Promega V5113) was then added at a ratio of 10ng trypsin per 1ug protein (no less than 500ng was added to a sample). The trypsin digestion was carried out at 37degC for 12–16 hours.

To stop the trypsin, formic acid (Fisher A117–50) was added at a ratio of 3uL per 100uL of sample to bring the pH down to less than 3. Peptides were cleaned up using an Oasis HLB uElution plate (Waters), equilibrated and washed with 0.04% trifluoroacetic acid in water, and eluted in 80% acetonitrile with 0.2% formic acid. All solutions used are HPLC grade. Samples were then lyophilized.

To remove any variance produced by phosphorylated peptides, the samples were phosphatase treated. Peptides were resolubilized in 50uL of 1X NEBuffer 3 (no BSA), and calf intestinal alkaline phosphatase (NEB M0290S) was added at a ratio 0.25 units per ug of peptide and incubated for 1 hour at 37degC. The peptides were cleaned up again according to steps described above. Peptides were resolubilized in 2% acetonitrile and 0.1% formic acid before SRM analysis.

### SRM Data Acquisition

As detailed in previous publications (Abell et al., 2011; Ahrends et al., 2014), 2μg of peptides were separated on an EASY-nLC Nano-HPLC system (Proxeon, Odense, Denmark) with a 200 mm × 0.075 mm diameter reverse-phase C18 capillary column (Maisch C18, 3 μm, 120 Å) and were subjected to a linear gradient from 8 to 40% acetonitrile over 70 min at a flow rate of 300 nl/min. Peptides were introduced into a TSQ Vantage triple quadrupole mass spectrometer (Thermo Fisher Scientific, Bremen, Germany) via a Proxeon nanospray ionization source. The transitions for the light (endogeneous) and heavy (SpikeTide) peptides were measured using scheduled SRM-MS and analyzed using Skyline version 3.5 (MacCoss Lab, University of Washington). Relative peptide quantifications were determined by ratioing the peak area sums of the transitions of the corresponding light and heavy peptides. Only transitions common between the heavy and light peptides with relative areas that were consistent across all samples were included in the quantification. Lists of transitions used for the 25-egg measurements in Figures 1F, 2A, and 2D and for the 120-egg measurements in Figures 2B and 3A are given in Tables EV3 and EV4, respectively.

### SRM Data Statistical Analysis

To minimize sample processing differences, a maximum of 30 single eggs were prepped and analyzed at the same time by SRM mass spectrometry. While we normalized the amount of heavy reference peptides added to each egg extract to the measured single egg protein concentration, this leaves still a small measurement error between individual eggs. This is likely both a result of small errors in the measurement of protein concentration and small volume pipetting errors, causing small under- or overestimation of relative protein abundances in a sample. This small calibration error was in previous protocols corrected using a normalization factor measured as a median of a set of anchor protein peptides (Abell et al., 2011; Feng and Picotti, 2016; Ludwig et al., 2012). Here we used the median of 22 normalized peptide intensities that minimally change during the cell cycle to derive a concentration correction factor for each egg (this factor was typically between 0.9 and 1.1). The lack of change in expression of these proteins during the cell cycle can be seen in Figure 1D. The correction we used makes the assumption that the 22 peptides are not overall co-regulated in the same direction, an assumption that is supported by both our SRM-MS and immunohistochemistry experiments (Figure 4). Specifically, we measured for a set of analyzed single eggs (e.g. 25 eggs in Figure 2A) the medians of the relative abundances for each of the 22 peptides across all eggs. To obtain a correction factor for each egg, we first normalized each peptide by the median of that particular peptide across all samples of interest (for example, for the 25-egg analysis show in Figure 1D, each peptide value was first divided by the median of that peptide across all 25 samples). Then we calculated the median of the 22 normalized peptide values for each egg. The resulting correction factor value was typically in the range of 0.9 to 1.1, and we divided all 26 relative protein abundances from that egg by this factor. The variation and co-variation values shown in this paper use these corrected relative abundances.

### Cell culture

MCF10A cells (ATCC, CRL-10317) were cultured in a growth media consisting of DMEM/F12 (Invitrogen) supplemented with 5% horse serum, 20 *μ*g/ml EGF, 10 *μ*g/ml insulin, 0.5 *μ*g/ml hydrocortisone, 100 ng/ml cholera toxin, 50 U/ml penicillin, and 50 *μ*g/ml streptomycin. HeLa cells were cultured in DMEM (Invitrogen) plus 10% fetal bovine serum (FBS) and penicillin-streptomycin-glutamine (PSG).

### EKAR-EV-NLS stable cell line

pPBbsr2-EKAR-EV-NLS was described previously (Komatsu et al., 2011). To generate stable cell lines, the construct was co-transfected with the piggybac transposase vector using polyethylenimine (Yusa et al., 2009). Cells with stable integration of the vector were selected for using 10ug/ml blasticidin (Invivogen).

### Immunofluorescence

Cells were fixed by adding paraformaldehyde to the cell media for 15 minutes (final concentration of paraformaldehyde in media was 4%). Cells were then washed three times in PBS before they were permeabilized by adding 0.2% triton X-100 for 20 minutes at 4 degC before being washed again with PBS. To remove cell size effects, cells were then stained with Alexa 647 NHS Ester as a marker of total protein mass and surrogate for cell volume/thickness following protocols described in (Kafri et al., 2013). The Alexa 647 NHS Ester was added at a concentration of 0.04 ug/mL in PBS for 1 hour. After washing again in PBS, a blocking buffer consisting of 10% FBS, 1% BSA, 0.1% triton X-100, and 0.01%NaN_3_ in PBS was added, and the cells were incubated for 1 hour at room temperature. Then primary antibodies were added overnight at 4 degC, followed by incubation with secondary antibodies for 1 hour at room temperature. To obtain particular protein concentrations for each cell, the mean total cell intensities of the respective antibodies were ratioed over the mean total cell intensity of the Alexa 647 NHS Ester.

### siRNA transfection

siRNAs were used at a final concentration of 20nM and are listed in the Key Resources Table (Table EV5). MCF10A and Hela cells were reverse-transfected with siRNA using Lipofectamine RNAiMax according to the manufacturer’s instructions. The cells were fixed 48 hours after reverse transfection with siRNA.

### Image Acquisition

For both fixed and live-cell imaging, cells were plated in 96-well, optically clear, polystyrene plates (Costar #3904). Approximately 10,000 HeLa cells or 5000 MCF10A cells were plated per well. For MCF10A cells, the wells were first coated with collagen (Advanced BioMatrix Cat #5005, PureCol Type I Bovine Collagen Solution) by placing fifty ul of collagen dissolved at a ratio of 1:100 in PBS in each well, incubating for 2–3h at room temperature, and then rinsing 3 times with PBS. MCF10A cells were then plated into the wells in MCF10A growth media. For assays to determine EGF responses, the media was aspirated from the cells 24 hours after plating and replaced with serum starvation media for 60 hours (DMEM/F12, 0.3% BSA, 0.5ng/mL hydrocortisone, 100ng/ml cholera toxin, PSG). For imaging, the cells were placed into an extracellular buffer consisting of 50mM KCl, 1.25M NaCl, 200mM Hepes, 15 mM MgCl2, 15 mM CaCl2 and 10 mM glucose. Time-lapse imaging was performed initially in 75 ul of extracellular buffer per well to which an additional 75ul of extracellular buffer containing 2X EGF doses was added to stimulate the cells.

Cells were imaged in a humidified 37degC chamber at 5% CO2. Images were taken every 2 min in the CFP and YFP channels using a fully automated widefield fluorescence microscope system (Intelligent Imaging Innovations, 3i), built around a Nikon Ti-E stand, equipped with Nikon 20x/0.75 N.A. objective, an epifluorescence light source (Xcite Exacte), and an sCMOS cameras (Hammatsu Flash 4), enclosed by an environmental chamber (Haison), and controlled by SlideBook software (3i). Five non-overlapping images were taken per well.

### Image Processing and Analysis

#### Segmentation and Tracking

Cell segmentation and tracking were performed using the “MACKtrack” package for MATLAB available at http://github.com/brookstaylorjr/MACKtrack, and described in (Selimkhanov et al., 2014). In place of the first-step cellular identification using differential-interference-microscopy, the first pass whole-cell segmentation was performed here by thresholding the total protein stain image.

#### Signal Measurement

Four channel fluorescent images were taken with a 10x objective on a MicroXL microscope, and image analysis was performed using Matlab analysis. Background subtraction was used in the Hoechst (to stain DNA and mask the nucleus), the two immunofluorescence, and the protein mass fluorescent channels. Signal intensities were corrected for non-uniformity but were still restricted to a central R=350 region of 2×2 binned images (1080×1080 pixels) of the image to minimize potential spatial non-uniformities in illumination and light collection towards the corners. The Hoechst stain was used to establish a nuclear mask and to select cells in the 2N G0/G1 state based on the integrated DNA stain. The Hoescht intensity levels used to define cells in the 2N state were selected by inspection of the Hoescht histograms. The live-cell FRET measurements of nuclear ERK activity were performed on a Nikon Ti2 controlled by 3i software (Intelligent Imaging, Denver, CO). The mean nuclear intensities of the FRET and CFP channels were ratioed for each cell to obtain the normalized FRET value at each timepoint. At the end of the timecourses, the cells were fixed and stained with either an ERK or MEK antibody, as well an Alexa 647 NHS Ester as an estimate of cell volume. To obtain ERK and MEK concentrations for each cell, the mean total cell intensities of the ERK and MEK antibodies were ratioed over the mean total cell intensity of the Alexa 647 NHS Ester. The final ERK and MEK concentrations for each cell were then matched to the corresponding FRET timecourse for that particular cell.

### Description of simulations of the multistep signaling pathway in Figures 1, 5, and 8

We used Matlab simulations of analog and binary signal transmission to help explain the consequence of expression variation in the concentration of pathway components and covariation between the components. We used a 5-step linear signaling pathway with a single input and output as an example of a typical vertebrate signaling pathway (see below for precise assumptions and calculation). The model was not saturated and uses a single fold-input R to increase pathway activation linearly above the basal activity level. The last regulated signaling step y5 is shown as the analog output A* in Figure 1. We simulated protein expression variation of each of the 10 signaling pathway components using lognormal Monte Carlo noise simulations (we multiplied each of the 10 system parameters by randomly variable factors centered on 1). We followed the system over time using the ODE45 function until it reached equilibrium at t=15.

We simulated binary pathways in Figure 5 by adding to the model an assumption that cells trigger a binary switch when the exceed a threshold level of 10 in the output y5*. Instead of plotting the distribution of the output level y5*, we plotted the fraction of cells that had an output level greater than 10 (fraction of cells in the active B* state). We used increasing fold-stimuli strength R and analyzed in the simulations the increasing fraction of cells that triggers the switch (plotted in the panels as a fraction of cells in a population activated versus strength of input stimulus R).

We also compared uncorrelated variation versus correlated variation (covariation) between signaling components in the pathway. In Figure 1E, we made the assumption that the 5 positive elements and the 5 negative elements in the model (possibly reflecting protein kinases versus protein phosphatases) each have a correlated variation. We compare this to the case were all variations are independent of each other as we also do in all other Figure panels. This correlated variation leads to an increase of the overall variation of the signaling response of a cell.

#### Linear model

The 10 lognormal stochastic values of a factorial parameter e(1-10) are calculated for each of typically 5000 runs to generate the plots, e(i:10)=(exp(randn(10,1), Var) in Matlab. Var is the %variation parameter that changes in different panels in the plots. Calculating a coefficient of variation (CV) of the resulting random parameter distribution returns the value Var.

A* corresponds to y5* and denote the final output signal. For each step, we assumed that the active y* states are generated from a relatively larger constant pool of precursors y1, y2, y3, y4 and y5 (the model is not saturated).

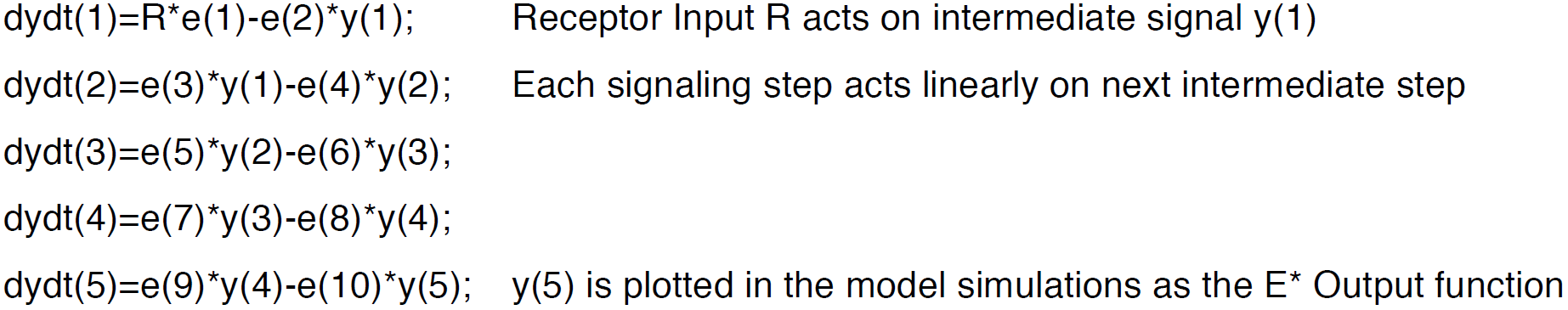

The calculations of fIDL and aHC were done analytically using the inverse normal distribution function to determine the fraction of cells in a population that extend past a given limit. In the case of the analog fold-Input Detection Limit, the value is half of the final signal output amplitude since both the unstimulated and stimulated distributions are symmetrical when they are plotted as a log scaled distribution see Figure 1C (black and green distributions). When assuming 95% accuracy for distinguishing stimulated and unstimulated cells from the output signal, this results in a fold-Input Detection Limit of:

fIDL = exp(CV* sqrt(N)*norminv(0.95)*2).

If the system has co-variation in components of the pathway, error propagation leads to a term sqrt(N-Ncov + Ncov^2) instead of sqrt(N), with Ncov being the number of co-varying components and N being the number of total components.

Figure EV1 compares fIDL values to the log2 Mutual Information content of the same system (bits), adding different levels of saturation to the last term of the equation (as an example, we used instead of y(4) the term 10*y(4)/(y(4)+9) for the system that imposes a saturation of a factor of 10 to the output signal). For the mutual information calculations, 10,000 simulations were made with R values spread out using a random number generator in log2 units. Output A* (log2 units) were simulated and the MI was derived from R and A* by using log2 in the MI equation and by using binning of 0.05 for R and A*.

A similar analysis can also be made to determine a Hill coefficient that fits the fraction of activated cells plotted as a function of the fold-input stimulus in Figure 5. The Hill coefficient for a given binary threshold Thr, can be derived from aHC=log(Thr)/log(R0), with R0 derived from the equation R0=exp(CV*sqrt(N)*norminv(Thr/(Thr+1)). This results in an:

HC = log(Thr)/(norminv(Thr/(Thr+1))*CV*sqrt(N))

For broad ranges of chosen Threshold values Thr for the binary switch, this equation is independent of the actual Threshold value used as long as it is significantly larger than the output distribution width for unstimulated cells.

In Figures 8B-F, we are combining these two curves by multiplying the logarithm of fIDL with HC to show they co-dependency:
log(fIDL)*HC = log(Thr)/norminv(Thr/(1+Thr))*norminv(0.95) ~ 6.3

### Description of ERK-MEK model in Figure 6

For numerical simulations, we used the ODE model of the ERK signaling network from (Sturm et al., 2010) with negative feedback intact. The model incorporates dynamics from RasGTP through Raf and MEK down to ERK phosphorylation. We used the input concentration of RasGTP as a proxy for extracellular EGF. The output was defined as doubly-phosphorylated ERK (pERK), which serves as a proxy for ERK activity, as ERK activity is a monotonically increasing function with respect to pERK. All model and simulation files can be found at http://github.com/TeruelLab/ERKmodel_v1.0.

## Acknowledgements

This work was supported by National Institutes of Health RO1-DK101743, RO1-DK106241, and P50-GM107615 (to M.N.T.), Stanford BioX Seed Grant funding (to M.N.T.), and T32-NIH T2HG00044 (to M.L.Z.). We thank James Ferrell, Tobias Meyer, and members of the Teruel Lab for discussions and critical reading of the manuscript.

## Author Contributions

K.M.K, M.L.Z., B.T. and M.N.T. conceived experiments. K.M.K. and B.T. performed experiments. K.M.K., B.T., and M.N.T. carried out model simulations. K.M.K, M.L.Z., B.T. and M.N.T. analyzed data. K.M.K. and M.N.T wrote the paper with input from all authors.

## Conflict of interest

The authors declare that they have no conflict of interest.

## LEGENDS FOR EXTENDED VIEW FIGURES

**Figure EV1. Comparison of fIDL and Mutual Information (MI) analysis.** MI analysis requires a fold-output range which we added to the model in Figure 1A by using a saturation term for y4 (see Methods). As shown for Expression Variations of 5 and 10 %, in contrast to MI analysis (top), fIDL analysis (bottom) is largely independent of the fold-output range.

**Figure EV2. Bootstrap analysis of CVs of the relative abundance of 26 proteins using a 60-egg set collected at 60 minutes after egg activation.** Bootstrapping of random samples was performed 2000 times with replacement. We used the bootstrap analysis to determine CVs for the entire 60-egg data set (blue circles) or on 6 batches of 10 eggs that were sequentially analyzed on the mass spectrometer (red circles). The lower CVs for batches of sequentially analyzed cells (median CV of 9% for the 26 proteins) argues that accurate concentration comparison using SRM analysis is optimally performed in batches of samples analyzed sequentially.

**Figure EV3. Comparison of technical and biological variation in the SRM mass spectrometry measurements.** To measure technical variability, 30 individual eggs were lysed and mixed together to collapse any biological variability. The lysate mixture was then pipetted into 30 individual tubes and processed individually before SRM analysis to quantify sample handling variation. CVs were compared to CVs measured in Figure 3B.

**Figure EV4. Representative images of immunostaining of the proteins studied.**

(A) Images from HeLa cells.

(B) Images from MCF10A cells.

**Figure EV5. siRNA-mediated depletion experiments to validate specificity of the respective antibodies and to test for co-regulation of protein expression of (A) MCM5/MC7M7 and (B) MEK/ERK.** HeLa cells were transfected with the respective siRNA. Forty-eight hours later, the cells were fixed, stained, and imaged to quantify expression of the respective proteins. Approximately 1000 cells were quantified in each histogram and barplot. Error bars show s.e.m. Results shown are representative of 3 independent experiments.

